# Interactions between *Pseudomonas aeruginosa* and six opportunistic pathogens cover a broad spectrum from mutualism to antagonism

**DOI:** 10.1101/2024.03.22.586229

**Authors:** Clémentine Laffont, Tobias Wechsler, Rolf Kümmerli

## Abstract

Bacterial infections often involve more than one pathogen. While it is known that polymicrobial infections can impact disease outcomes, we have a poor understanding about how pathogens affect each other’s behaviour and fitness. Here, we used a microscopy approach to explore interactions between *Pseudomonas aeruginosa* and six opportunistic human pathogens that often co-occur in polymicrobial infections: *Acinetobacter baumannii*, *Burkholderia cenocepacia*, *Escherichia coli*, *Enterococcus faecium*, *Klebsiella pneumoniae*, and *Staphylococcus aureus.* When following growing micro-colonies on agarose pads over time, we observed a broad spectrum of species-specific ecological interactions, ranging from mutualism to antagonism. For example, *P. aeruginosa* engaged in a mutually beneficial interaction with *E. faecium* but suffered from antagonism by *E. coli* and *K. pneumoniae*. While we found little evidence for active directional growth towards or away from cohabitants, we observed that certain species increased growth in double layers in co-cultures and that physical forces due to fast colony expansion had a major impact on fitness and interaction patterns. Overall, our work provides an atlas of pathogen interactions, potentially useful to understand species dynamics in polymicrobial infections. We discuss possible mechanisms driving pathogen interactions and offer predictions of how the different ecological interactions could affect virulence.

## Introduction

*Pseudomonas aeruginosa* is an ubiquitous bacterium, widespread in natural environments such as water and soil, with a high occurrence in areas that are exposed to human activities (Crone et al., 2020; Green et al., 1974; Mena and Gerba, 2009). It is also one of the major opportunistic human pathogens responsible for nosocomial infections. It causes serious complications in patients suffering from cystic fibrosis, and further causes pneumonia, burn wound infections, necrotizing skin, bloodstream infections, and perichondritis in immuno-compromised patients (Costa et al., 2015; de Bentzmann and Plésiat, 2011; Mulcahy et al., 2014; Murray et al., 2007; Reynolds and Kollef, 2021; Spernovasilis et al., 2021). In addition, infections with this bacterium can involve multiple other pathogens, and such polymicrobial infections are often associated with higher host morbidity (Bisht et al., 2020; Cendra and Torrents, 2021; Murray et al., 2014; Rogers et al., 2003). Polymicrobial infections are increasingly considered as prevalent (Azevedo et al., 2017; Brogden et al., 2005; Peters et al., 2012) and of major concern because interactions between pathogens often complicates treatment options (Al-Wrafy et al., 2023; Filkins and O’Toole, 2015; Rocha-Granados et al., 2020).

Studying pathogen interactions has become a prospering field (Dunny et al., 2008; Kramer et al., 2020; Michie et al., 2016; Ross and Whiteley, 2020; West et al., 2007a). Especially, interactions between key pathogens like *P. aeruginosa* and *Staphylococcus aureus* have been extensively studied as these species often co-occur in patients (Hotterbeekx et al., 2017; Ibberson and Whiteley, 2020; Limoli and Hoffman, 2019; Nguyen and Oglesby-Sherrouse, 2016). Research focused on various aspects including molecular mechanisms of interactions (Armbruster et al., 2016; Limoli et al., 2019; Yarrington et al., 2024; Zarrella and Khare, 2022), variation in interactions between different strains and across environments (Bernardy et al., 2022; Niggli et al., 2021; Niggli and Kümmerli, 2020), the evolution of interactions (Niggli et al., 2023; Tognon et al., 2017) and the consequences of interactions for the host (Orazi and O’Toole, 2017; Radlinski et al., 2017; Rezzoagli et al., 2020). Interactions among other pathogens (e.g. *Klebsiella pneumoniae* vs. *Escherichia coli* and *P. aeruginosa* vs. *Burkholderia cenocepacia*) have also been studied (Chattoraj et al., 2010; Juarez and Galván, 2018; Leinweber et al., 2018; Morin et al., 2022), but to a lesser extent. A general insight from this body of work is that competition [-/-] and antagonism [+/-] seem to prevail (Schmitz et al., 2023), although rarer cases of mutual benefits [+/+] have also been reported (Camus et al., 2020). Despite these insights, we know still little about the generality of interaction patterns. For example, does a specific pathogen show standard responses to all other pathogens, or are interaction patterns specific to the identity of the cohabitant? Moreover, are interactions indeed dominated by competitive and antagonistic interactions or are there also opportunities for more neutral interactions, such as neutralism [0/0], commensalism [+/0] or amensalism [-/0], where at least one species is not affected by the presence of the other one (Figure 1A)?

**Figure 1:**
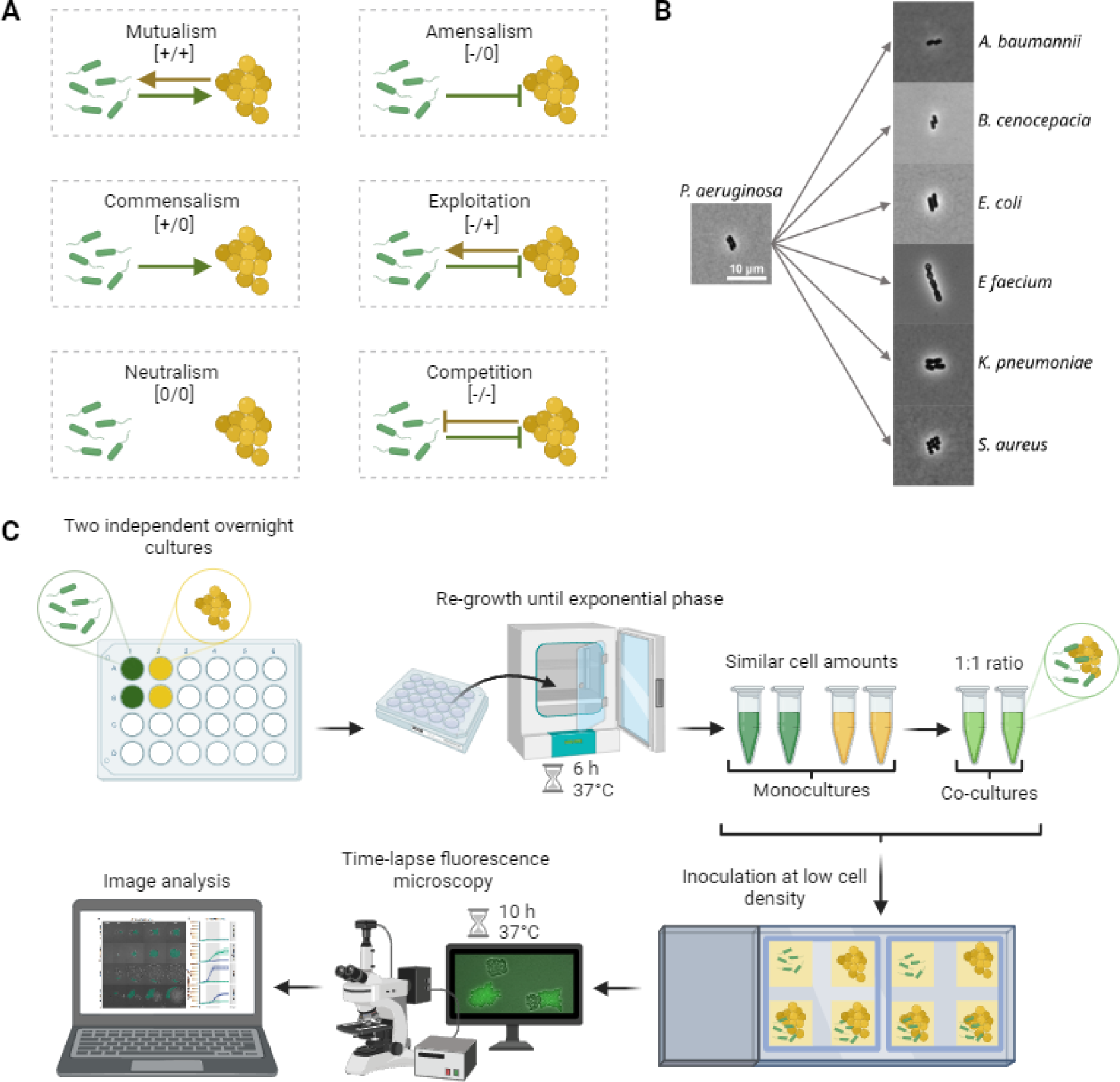
Scheme of agarose-pad assay for the study of interactions between *P. aeruginosa* and six other pathogens. A) Spectrum of ecological interactions found in microbial communities. The arrows and stop arrows respectively show positive and negative impact of one species on another. B) Snapshots of cells of the seven pathogen species used in the interaction assays. The scale of all pictures is 10 µm. C) Workflow of the interaction assay. After adjusting the optical density (OD at 600nm) from overnight cultures, *P. aeruginosa* and the cohabitant species were re-grown until the exponential phase. ODs were then adjusted once more to obtain similar cell numbers. Mono- and co-cultures (1:1 ratio) were inoculated at low cell density on agarose pads. Time-lapse fluorescence microscopy was carried out for 10 hours at 37°C with pictures taken every five minutes. After a drift-correction and image cropping, colonies were segmented automatically. *P. aeruginosa* featured a constitutively expressed GFP marker do distinguish its colonies from colonies of the cohabitant species. The area, the shape and the growth directionality of the colonies were measured over time using automated scripts in ImageJ and Rstudio. This figure was created with BioRender.com.

Here, we aim to address these questions by using *P. aeruginosa* as our focal pathogen in co-cultures with six other pathogenic species that often co-occur with *P. aeruginosa* in polymicrobial infections: *Acinetobacter baumannii*, *B. cenocepacia*, *E. coli*, *Enterococcus faecium*, *K. pneumoniae*, and *S. aureus* (Cendra and Torrents, 2021; Françoise and Héry-Arnaud, 2020; Gaston et al., 2021; Heitkamp et al., 2018). We conducted our experiments on agarose pads, where we tracked the growth and interactions of species from the single-cell stage to the microcolony level using time-lapse fluorescence microscopy (Figure 1B-C). Agarose pads provide a structured environment that mimic more closely (as compared to liquid medium) the surface-attached mode of growth pathogens follow in infections (Bjarnsholt et al., 2013; Donlan, 2002). Thanks to the time-lapse approach, we can quantify the fitness of each species in mono- and co-cultures (Limoli et al., 2019; Niggli et al., 2021) and thereby determine the type of interaction occurring between pathogens (Figure 1). Furthermore, our approach allows us to quantify changes in bacterial behaviour over time in expanding colonies. Specifically, we can quantify whether (i) pathogens show directional growth towards or away from each other, (ii) physical forces due to colony expansion affect colony morphology, and (iii) there are any other form of behavioural changes. Finally, while our assay purely tracks behaviour and fitness over time, we offer a detailed discussion on the putative mechanisms involved in pathogen-pathogen interactions, generating workable hypotheses for future molecular studies.

## Results

### Growth and colony morphology highly vary across pathogens

In a first assay, we tracked colony morphology (Figure 2A) and growth (Figure 2B) of all seven bacterial pathogens in monocultures on agarose pads. We observed that all species were able to grow, mainly between the 2^nd^ and 7^th^ hour. Growth, measured as the area of the colony, varied substantially between species (Figure 2B). We noticed relatively low growth for *A. baumannii* and *B. cenocepacia*, intermediate growth for *P. aeruginosa* and *S. aureus*, and high growth for *E. coli*, *E. faecium* and *K. pneumoniae*. We further observed differences in colony morphologies (Figure 2A). Colony shapes varied from elongated for *P. aeruginosa*, to roundish for *A. baumannii*, *B. cenocepacia* and *E.coli,* to circular for *E. faecium*, *K. pneumoniae* and *S. aureus*.

**Figure 2:**
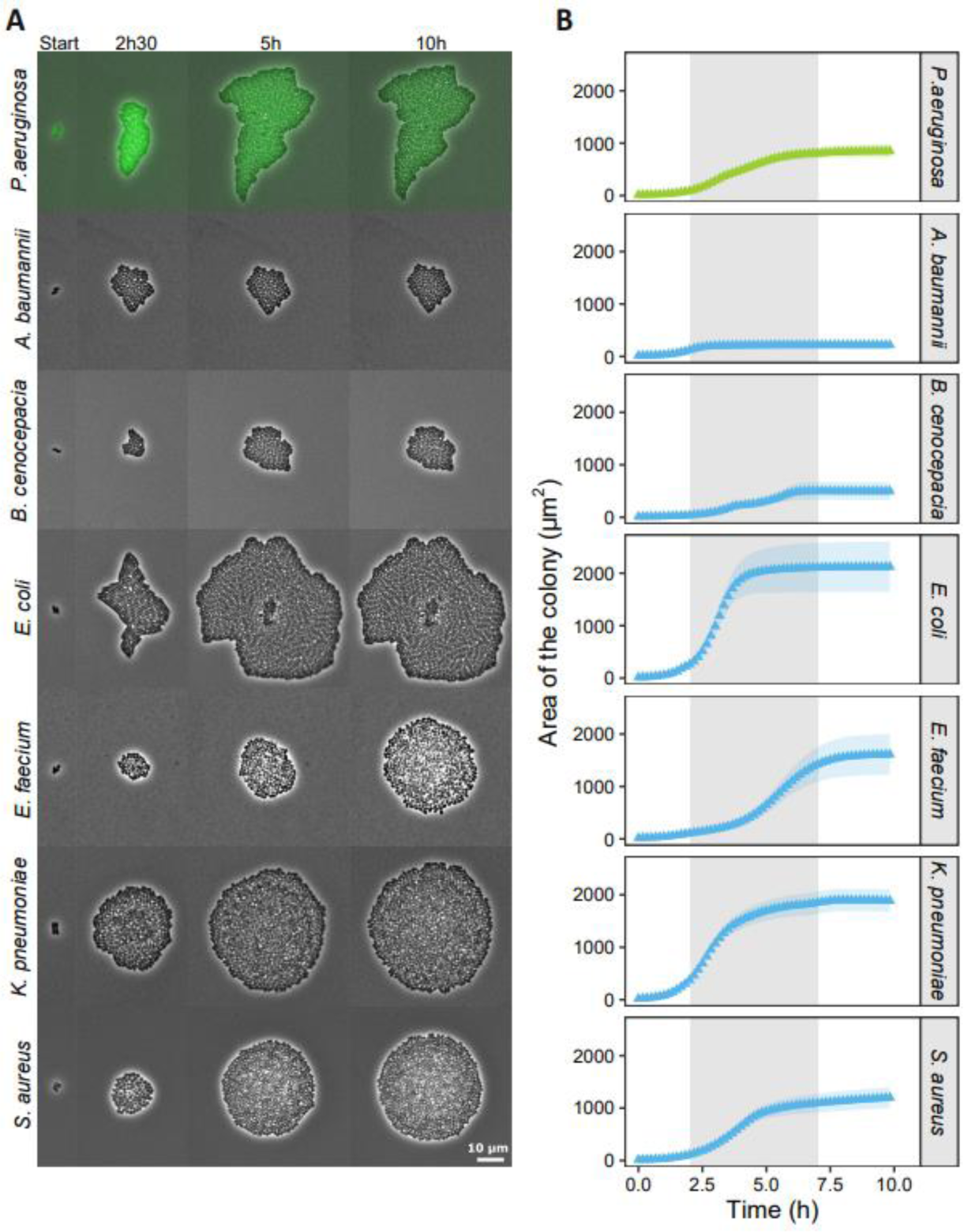
Monoculture growth and colony morphology of the seven bacterial pathogen species. (A) Snapshots of representative colonies of the different species grown on agarose pads. *P. aeruginosa* (in green) is tagged with a constitutively expressed GFP marker. Time-lapse pictures are shown for four time points (indicated on top of the panels). The contrast of the pictures was enhanced to improve colony visibility. (B) Colony growth curves of the seven pathogen species based on the area occupied by colonies. Data points and shaded areas represent the means and the 95% confidence intervals of colony growth, respectively. *P. aeruginosa* is shown in green, while the other species are shown in blue. The grey shaded area indicates the time window (2^nd^ to 7^th^ hour) during which most of the growth occurred.

### P. aeruginosa growth and colony morphology change in co-cultures with other pathogens

Next, we investigated the colony morphologies (Figure 3A) and growth (Figure 3B) of *P. aeruginosa* (tagged with a GFP marker) in co-cultures with the six other pathogens (referred to as cohabitants). In the Supporting Information, we provide representative movies (Movie S1-S6) for all combinations. As in monocultures, all the species grew predominantly between the 2^nd^ and 7^th^ hour. We noticed that *P. aeruginosa* growth was drastically affected by the identity of cohabitants. Growth was low in co-cultures with *A. baumannii*, *E. coli* and *K. pneumoniae*, intermediate in co-cultures with *E. faecium* and *S. aureus*, and high in co-cultures with *B. cenocepacia* (Figure 3B). We observed that colonies of the two species often came into contact, which induced morphological changes in *P. aeruginosa* colonies in a cohabitant-specific manner. Specifically, *P. aeruginosa* colonies became more roundish in co-cultures with *A. baumannii* and *B. cenocepacia,* but more elongated and sometimes even shapeless in co-cultures with *E. coli*, *E. faecium*, *K. pneumoniae*, and *S. aureus.* In contrast, colony morphology remained largely unchanged in all six cohabitant species.

**Figure 3:**
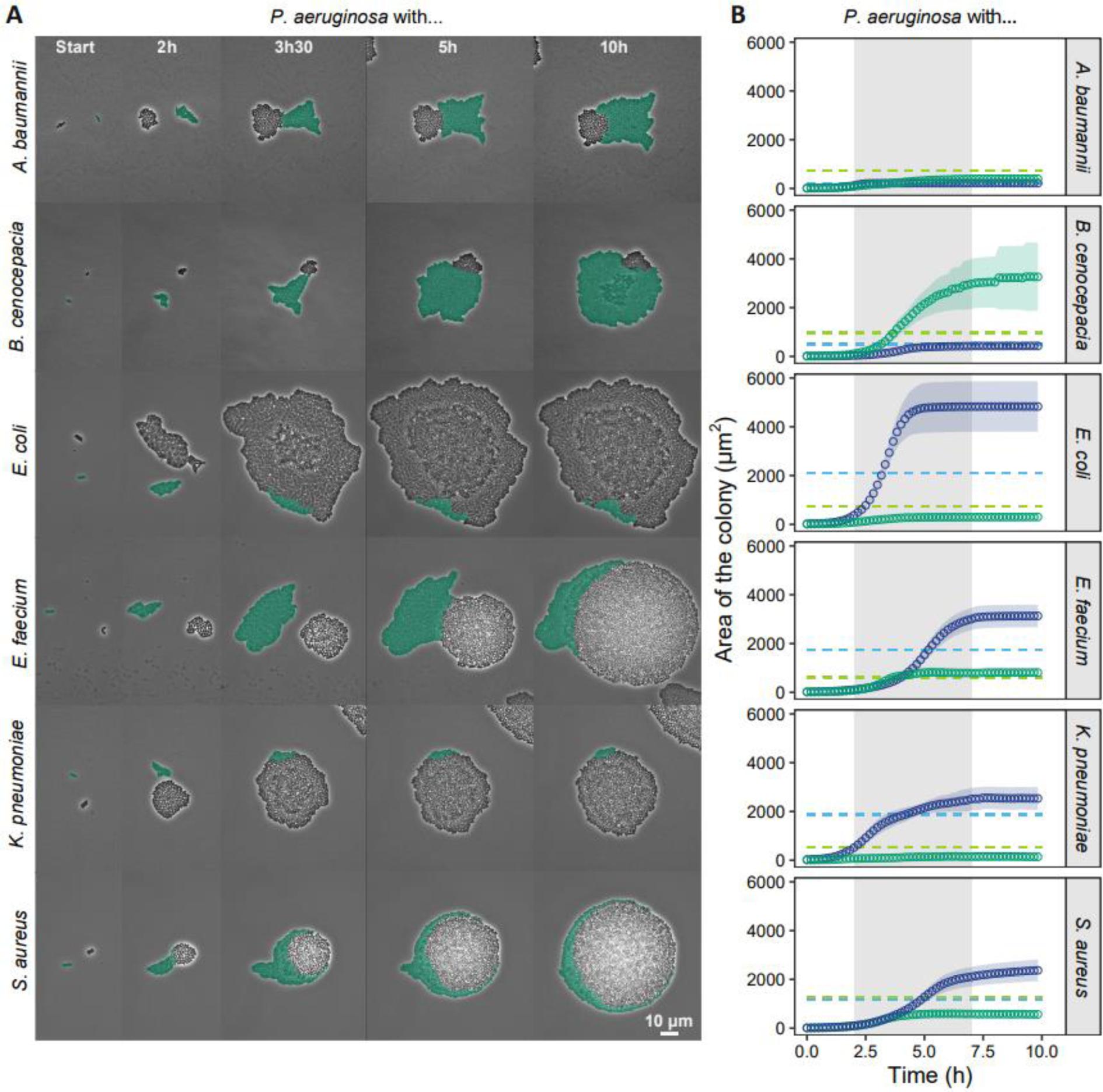
Growth and colony morphology of *P. aeruginosa* change in co-cultures with six different bacterial cohabitants. (A) Snapshots of representative colonies of the co-cultured species on agarose pads. *P. aeruginosa* (in green) is tagged with a constitutively expressed GFP marker (signal artificially coloured for illustration purposes). Time-lapse pictures are shown for five time points (indicated in the top panels). The contrast of the pictures was enhanced to improve colony visibility. Movies of the corresponding assays are available in the Supporting Information (Movies S1-S6). (B) Growth curves of the co-cultured species based on the area occupied by the respective colonies. Data points and shaded areas represent the means and the 95% confidence intervals of colony growth, respectively. *P. aeruginosa* is shown in green, while the cohabitants are shown in blue. The grey shaded area indicates the time window (2^nd^ to 7^th^ hour) during which most of the growth occurred. The light dashed lines show the mean of the growth yield of *P. aeruginosa* (green) and the cohabitants (blue) in monocultures.

### Diverse set of ecological interactions between P. aeruginosa and other pathogens

Following our descriptive assessment of pathogen growth on agarose pads, we quantified and compared the maximum growth rate (Figure S1 & Table S1) and the area under the growth curve (Figure 4 & Table S2) for each species combination in mono-versus co-cultures. The goal of this analysis is to statistically test whether pathogens affect each other’s fitness in a positive or negative way. Among the six *P. aeruginosa* / cohabitant combinations, our analysis yielded four different types of ecological interactions (Figure 4 & Table S2).

- Amensalism [-/0] occurred between *P. aeruginosa* and *A. baumannii* and between *P. aeruginosa* and *K. pneumoniae*. In both cases, *P. aeruginosa* experienced negative fitness effects in co-cultures (Tukey HSD *post-hoc* comparisons following two-way ANOVA: p = 0.0164 and p < 0.0001 respectively), while the fitness of the cohabitant was not affected (p = 0.8913 and p = 0.5960 respectively).
- Commensalism [+/0] occurred between *P. aeruginosa* and *B. cenocepacia*, whereby the fitness of *P. aeruginosa* increased in co-culture (p < 0.0001), while *B. cenocepacia* fitness was not affected (p = 0.9981).
- Mutualism [+/+] occurred between *P. aeruginosa* and *E. faecium,* as both species showed higher fitness in co-compared to monocultures (p = 0.0014 and p < 0.0001 respectively).
- Exploitation [-/+] occurred between *P. aeruginosa* and *E. coli* and between *P. aeruginosa* and
- *S. aureus*. In both cases, *P. aeruginosa* experienced negative fitness effects in co-cultures (p < 0.0001 and p = 0.0055 respectively), while the fitness of the other pathogens increased (p < 0.0001 and p = 0.0042 respectively).

**Figure 4:**
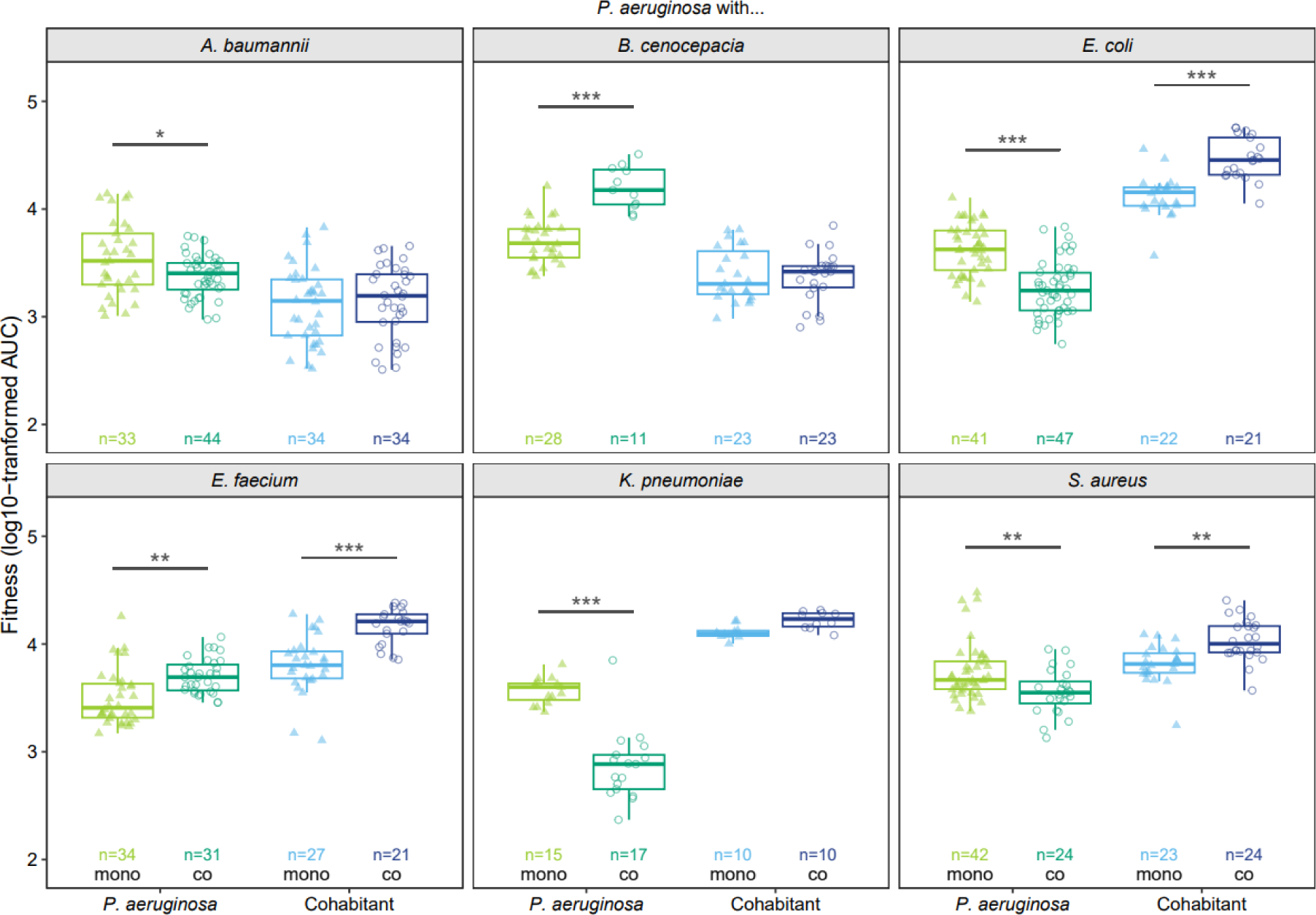
The fitness of at least one species is affected when *P. aeruginosa* is co-cultured with other pathogens. The boxplots depict the log10-transformed fitness (AUC: area under the colony growth curve) of *P. aeruginosa* (green) and its cohabitants (blue) in monocultures (light triangles) and co-cultures (dark circles). n-values indicate the total number of colonies tracked for each species combination. Two-way ANOVAs were used in combination with Tukey HSD *post-hoc* tests to examine fitness differences between mono- and co-cultures for *P. aeruginosa* and its cohabitants. Asterisks show the level of significance: * p < 0.05, ** p < 0.01, *** p < 0.001.

None of the interactions was neutral [0/0] or competitive [-/-] and in four out of the six combinations, at least one species experienced an absolute fitness benefit in co-cultures.

### Little evidence for increased directional growth in co-cultures

Given that many interactions were associated with positive or negative fitness consequences, we asked whether bacteria show increased levels of directional growth in co-compared to monocultures. Directional growth would allow bacteria to grow away or towards harmful or beneficial cohabitants, respectively. We measured the directionality for every hour separately, and then compared it between mono- and co-cultures during the time window of actual growth (2^nd^ to 7^th^ hour).

For *P. aeruginosa* monocultures, the growth directionality decreased over time (Figure 5). In contrast to our hypothesis, we found no changes in *P. aeruginosa* directionality between mono- and co-cultures in four out of six species combinations. In the remaining two combinations, *P. aeruginosa* growth directionally increased with *B. cenocepacia* and decreased with *E. coli*, but only at a few specific time points. Low levels of growth directionality change were also found for the six cohabitants (Figure 5). Growth directionality decreased or stayed stable over time in monocultures, and there were only very few instances where growth directionality significantly changed in mono-compared to co-cultures (increase for *B. cenocepacia* (one time point) and for *S. aureus* (two time points); decrease for *E. faecium* (one time point), Table S3). Taken together, pathogens show no or very moderate active movement away or towards co-growing species in our agarose-pad system.

**Figure 5:**
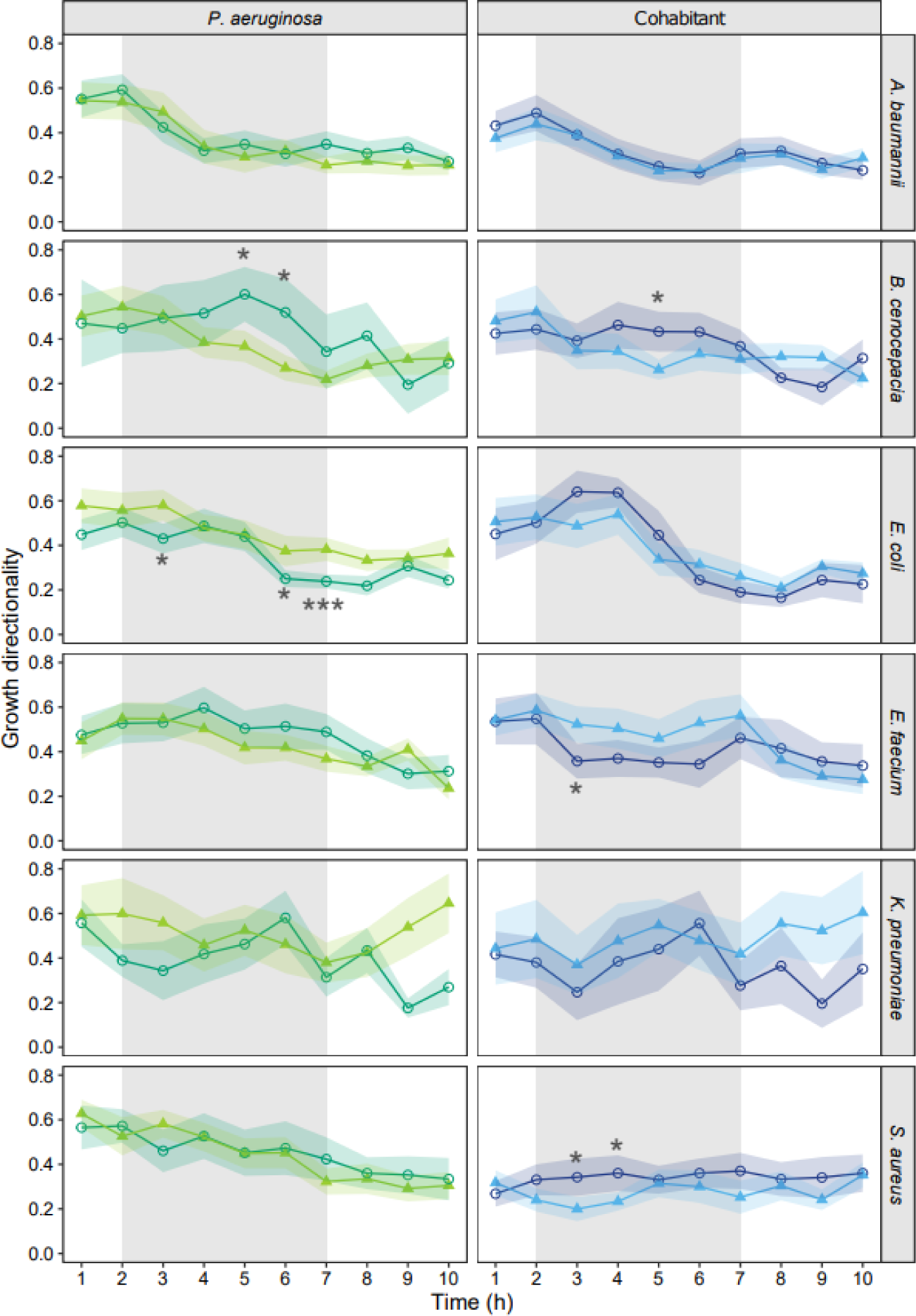
Colony growth directionality of *P. aeruginosa* and the six cohabitant species hardly differ between mono- and co-culture conditions. Growth directionality is calculated for each hour separately. Higher values stand for more directional growth. Data points and shaded areas represent mean values and 95% confidence intervals across colonies of *P. aeruginosa* (green) and its cohabitants (blue) for monocultures (light triangles) and co-cultures (dark circles). The grey shaded areas show the time window (2^nd^ to 7^th^ hour) during which most of the growth occurred. Welch’s two sample t-tests were used to compare the directionality between mono- and co-cultures, separately for each species, at each time point between the 2^nd^ and the 7^th^ hour. Asterisks show significance levels after FDR-corrections to account for multiple comparisons: * p < 0.05, ** p < 0.01, *** p < 0.001.

### Physical forces have a major impact on colony morphology

While the above analysis revealed little evidence for active behavioural changes in co-cultures, we asked whether passive forces could impact colony morphology. We hypothesize that colony morphology might passively change once colonies of the co-cultured species come into contact. We predict this effect to be particularly strong in co-cultures with the fast-growing cohabitants (*E. coli*, *E. faecium*, *K. pneumoniae* and *S. aureus*) because these species might simply push *P. aeruginosa* out of their way due to their fast colony expansion (Figure 6A). To track the effects of such physical forces, we quantified the roundness of colonies over time and predict that increased physical forces imposed by the co-growing species should decrease colony roundness.

**Figure 6:**
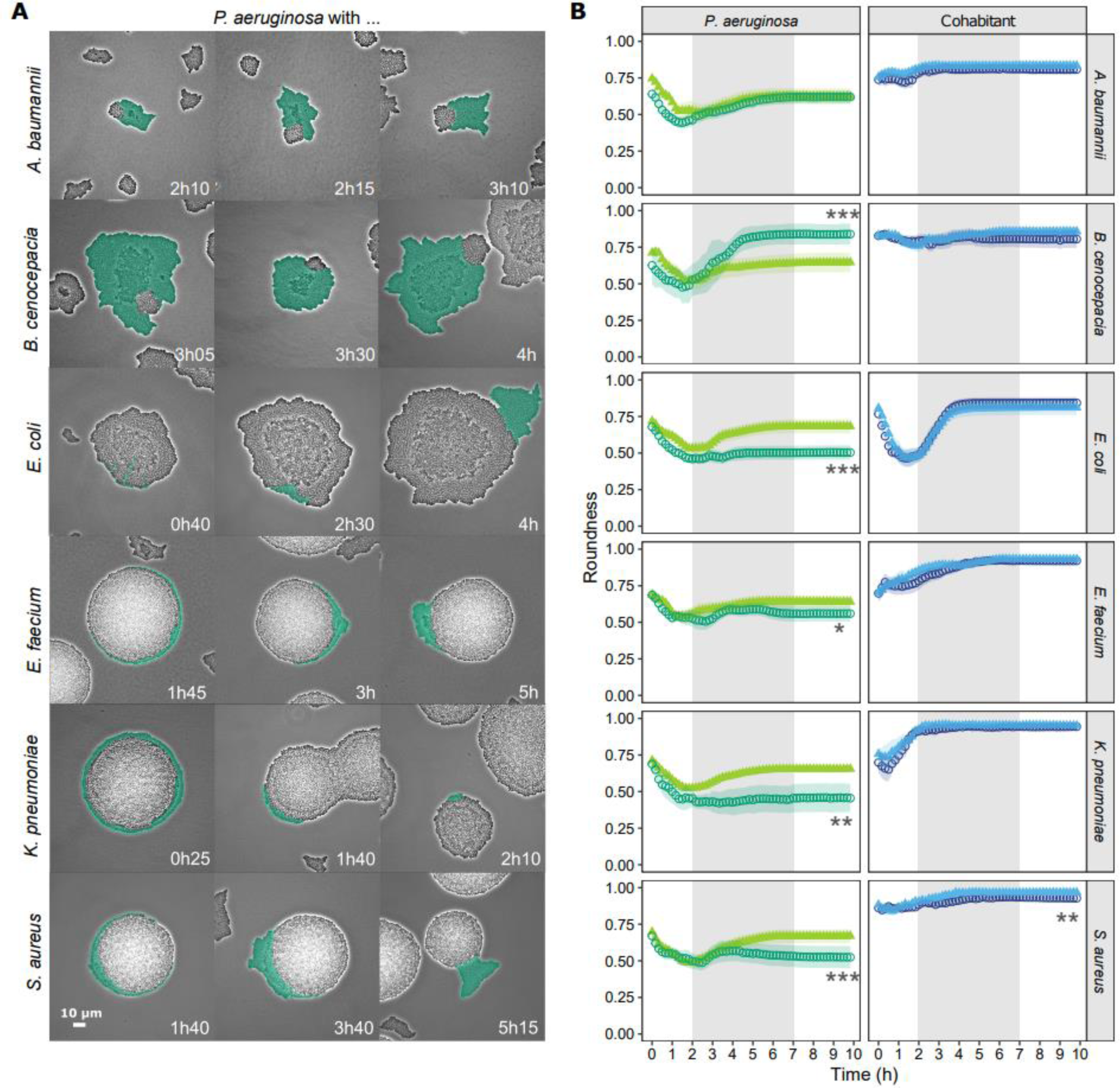
Physical forces due to rapid colony expansion of cohabitant species affect *P. aeruginosa* colony morphology. A) Snapshots of physical interactions between colonies of the co-cultured species. Three independent examples are shown for each species combination and the time points at which colonies came into contact are indicated in the bottom of each panel. *P. aeruginosa* (in green) is tagged with a constitutively expressed GFP marker (signal artificially coloured for illustration purposes). The contrast of the pictures was enhanced to improve colony visibility. B) Roundness of colonies over time for *P. aeruginosa* (green) and its cohabitants (blue) in monoculture (light triangles) and co-cultures (dark circles). Colony roundness can vary between one and zero, whereby values close to one indicate near-circular colonies. Data points and shaded areas show the mean values and the 95% confidence intervals across all colonies of the respective species and growth conditions. The grey shaded areas show the time window (2^nd^ to 7^th^ hour) during which most of the growth occurred. Welch’s two sample t-tests were used to compare the roundness between mono- and co-cultures, separately for each species, at the end of the experiment (10h). Asterisks show significance levels: * p < 0.05, ** p < 0.01, *** p< 0.001. For the experiments involving *K. pneumoniae*, we deviated from the standard procedure to estimate the roundness of *P. aeruginosa* for technical reasons (see methods).

For *P. aeruginosa*, we found strong support for this hypothesis (Figure 6B & Table S4). The roundness of *P. aeruginosa* colonies became significantly lower in co-compared to monocultures with the four fast-growing species (*E. coli*, p < 0.0001; *E. faecium*, p = 0.0194; *K. pneumoniae*, p = 0.0005; *S. aureus*, p = 0.0017), but only when contacts between colonies started to occur. Also compatible with our hypothesis, we observed that *P. aeruginosa* colony roundness significantly increased (p = 0.0002) in co-cultures with *B. cenocepacia* where *P. aeruginosa* itself is the faster growing species. In contrast to *P. aeruginosa*, we found that colony roundness was not different between mono- and co-cultures in five out of the six cohabitants, and only slightly dropped for *S. aureus* (p = 0.0001). Taken together, our results suggest that the plasticity in *P. aeruginosa* morphology is primarily driven by the passive physical forces exerted by (fast) growing co-cultured species, and not by active decisions taken by *P. aeruginosa*.

### Growth in double layers is increased in certain pathogen combinations

We manually screened time-lapse movies to detect any additional behavioural changes associated with pathogen growth in co-cultures. One effect we repeatedly observed in several species combination was the increased rate of growth in double layers in co-cultures compared to monocultures. We saw this effect for *P. aeruginosa* against *A. baumannii* (low extent) and *B. cenocepacia* (high extent), and for *E. coli* against *P. aeruginosa* (high extent) (Figure 3 and Figure 6). For *E faecium*, *K. pneumoniae* and *S. aureus*, double layering might also have occurred but was difficult to spot. Consequently, we quantified the double layer effect in the first three cases listed above. We found that both the frequency of and the area occupied by double layers was significantly increased in co-cultures (Figure 7 and Tables S5 + S6). In two of the three cases, increased double layer rates occurred in the species experiencing fitness benefits (*P. aeruginosa* growing with *B. cenocepacia* and *E. coli* growing with *P. aeruginosa*). Consequently, the double layer effect could be a consequence of increased growth combined with space limitations. Alternatively, it could also indicate that pathogens can detect cohabitants (possibly through diffusible molecules) and react to them in a species-specific manner.

**Figure 7:**
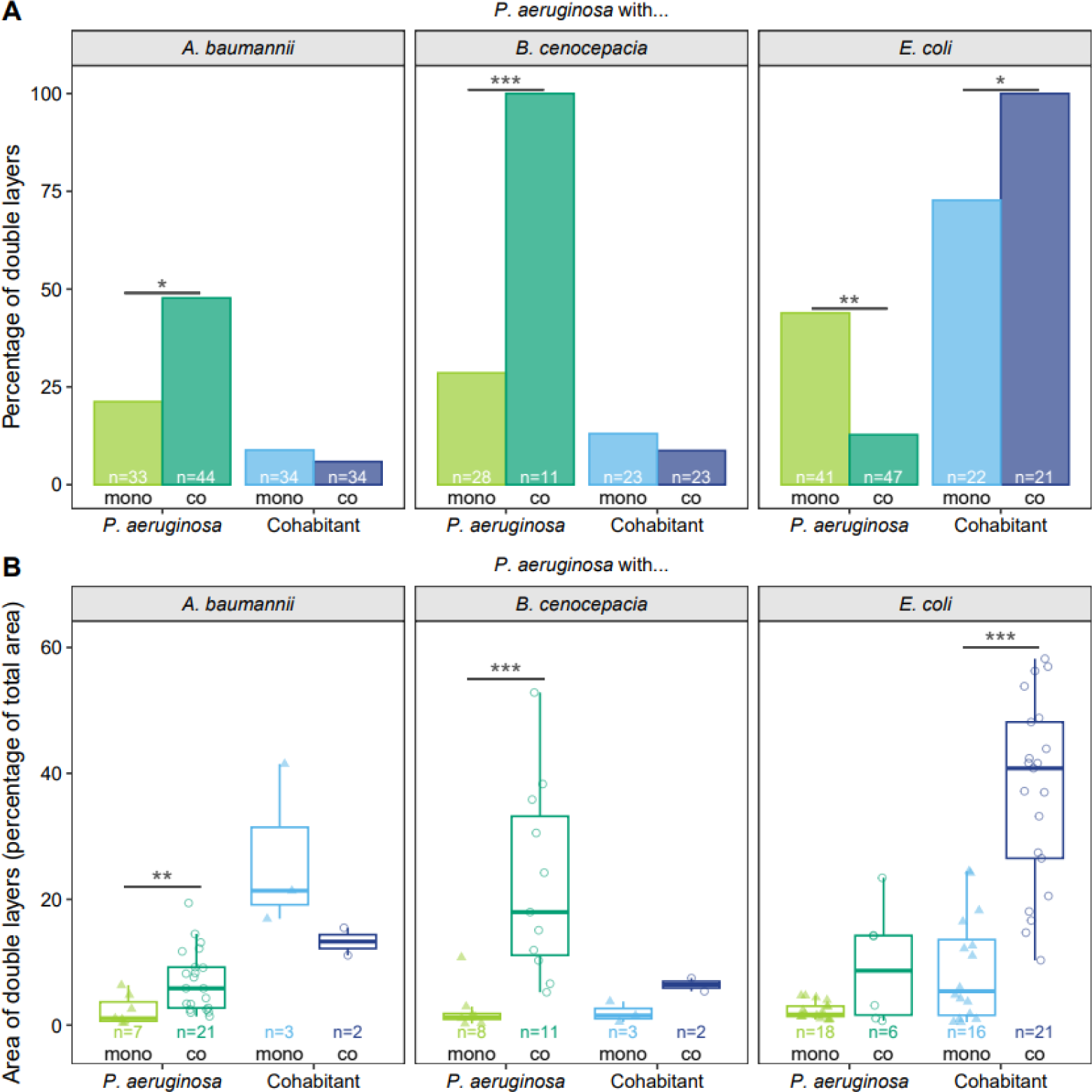
Certain pathogens respond to co-culturing by increased growth in double layers. A) Percentage of colonies of *P. aeruginosa* (green) and its cohabitants (blue) forming double layers in monocultures (light triangles) and co-cultures (dark circles). n-values indicate the total number of colonies tracked for each species combination. Fisher’s exact tests for count data were used to examine differences between mono- and co-cultures for *P. aeruginosa* and its cohabitants. B) Area of the double layers compared to the total area of *P. aeruginosa* (green) and its cohabitants (blue) in monocultures (light triangles) and co-cultures (dark circles). n-values indicate the total number of colonies tracked (number of colonies forming double layers) for each species combination. Welch’s two sample t-tests were used to examine differences between mono- and co-cultures for *P. aeruginosa* and its cohabitants. Asterisks show the level of significance: * p < 0.05, ** p < 0.01, *** p < 0.001.

## Discussion

The goal of our study was to measure fitness consequences and changes in bacterial behaviour of human opportunistic pathogens when cultured together on surfaces. We took *P. aeruginosa* as the focal species and cultured it together with six other pathogens that often co-occur with *P. aeruginosa* in infections. We measured species fitness by tracking colony growth using time-lapse fluorescence microscopy and observed four different ecological interactions among the six pathogen pairs, including amensalism [-/0], commensalism [+/0], mutualism [+/+] and exploitation [-/+]. This result indicates that pathogen-pathogen interactions are highly specific to the cohabitants involved and that a large pallet of ecological interactions is covered. In terms of behavioural changes, we found little evidence for directional movements of cells towards or away from cohabitants. However, we observed behavioural responses in terms of growth in colony double layers indicating that certain species react to cohabitants and change their growth patterns. Furthermore, we identified physical forces due to rapid colony expansion as a major force affecting colony morphology and fitness. Altogether, our simple assay visualizes and quantifies the complexity of pathogen-pathogen interactions in a spatially structured environment. Below, we discuss the potential effects of such interactions for the host and the putative mechanisms driving the observed fitness patterns.

A current debate in the field is whether mutualistic cooperation [+/+] or competition [-/-] prevail in interactions between microbes (Foster and Bell, 2012; Hibbing et al., 2010; Kost et al., 2023; Oliveira et al., 2014; Palmer and Foster, 2022; Piccardi et al., 2019; Rakoff-Nahoum et al., 2016; West et al., 2007b; Wingreen and Levin, 2006). Our results reveal a more nuanced picture by showing that in three out of six cases one of the two species experienced no change in fitness (two cases of amensalism and one case of commensalism). This suggests that many interactions might be accidental whereby the mechanism deployed by the neutral pathogen (*i.e*. experiencing no fitness consequences) might not have been selected for the purpose to harm or help others. However, it is important to differentiate between the ecological definition of social interactions (Figure 1A) and their evolutionary consequences. In evolutionary biology, competitive behaviours include those that increase the fitness of the actor relative to the cohabitant. This definition can principally apply to several ecological interactions – amensalism, exploitation, competition – as in all these cases the actor supresses the growth of cohabitants and can thereby earn relative fitness benefits. In this evolutionary context, four of the six interactions are competitive in our assay.

Understanding ecological interactions between pathogens could help to predict virulence outcomes in polymicrobial infections. A simple prediction is that interaction types involving negative fitness consequences (amensalism [-/0], competition [-/-]) should reduce virulence, whereas interaction types involving positive fitness consequences (commensalism [+/0], mutualism [+/+]) should increase virulence. These predictions are simplifications as they underlie the assumption that the mechanisms through which pathogens interact do not affect the host. This assumption can, for example, be violated by mechanisms that involve the secretion of broad-spectrum toxins that not only drive pathogen competition but may also adversely affect host tissue (O’Brien and Fothergill, 2017; Rada and Leto, 2013). Moreover, predictions will also depend on which pathogen is suppressed in interactions. Low or no reduction in virulence can be expected if the more virulent pathogen suppresses the less virulent pathogen in interactions. For example, a recent study revealed that *P. aeruginosa* is more virulent than *B. cenocepacia* in an insect host model in mono-infections. As *P. aeruginosa* also suppressed *B. cenocepacia* in the host, there was no overall change in virulence in co-infections with the two species (Schmitz et al., 2023). In another study, zebrafish larvae were infected with *P. aeruginosa* and either *A. baumannii* or *K. pneumoniae* (Schmitz et al., 2024). The latter two species are less virulent than *P. aeruginosa* but suppress it through amensalims according to our findings (Figure 4). In line with our prediction, virulence was reduced in co-infections as compared to *P. aeruginosa* mono-infections. Finally, several studies showed that mutualistic interactions between pathogens increase virulence (Harrison et al., 2006; Rezzoagli et al., 2020; Rumbaugh et al., 2009). While there is currently not enough data available to build strong predictive models, our considerations indicate that ecological interactions between pathogens may be good indicators of how host will be affected by specific species combination in polymicrobial infections.

While our work is entirely based on behavioural and fitness assays, we discuss four mechanistic factors that could be involved in driving the observed pathogen interactions. The first factor comprises physical forces imposed by colony expansion. We investigated this factor and found that species with rapid growth and colony expansion on surfaces (*E. coli*, *E. faecium*, *K. pneumoniae*, *S. aureus*) simply pushed *P. aeruginosa* colonies out of their way. This physical mechanism resulted in drastic deformation of *P. aeruginosa* colony shapes and was associated with negative fitness consequences for *P. aeruginosa* in three out of four cases (exception: co-cultures with *E. faecium*). While it is known that mechanistic forces affect colony morphology and expansion (Farrell et al., 2013; Grant et al., 2014; Lloyd and Allen, 2015), we here show that such forces can be imposed by co-growing species. The second factor involves beneficial or inhibitory compounds secreted by bacteria. There are numerous compounds including toxins (broad and narrow spectrum), enzymes (proteases, lipases), siderophores (iron-chelators), and metabolic products (amino acids) that bacteria actively or passively secrete (Germerodt et al., 2016; Kramer et al., 2020; Pierson and Pierson, 2010; Saising et al., 2012). We do not know to which extent these compounds played a role in our assays. However, it is not unrealistic to assume that the positive fitness benefits *P. aeruginosa* gained in co-culture with *B. cenocepacia* and *E. faecium* accrue due to secreted compounds. They may work in concert with the above-described physical forces, whereby the latter increase the contact surface between the two species, which in turn favours a more efficient compound uptake. The third factor includes contact-dependent killing and inhibition systems such as T6SS and CDI (Hernandez et al., 2020; Ikryannikova et al., 2020). Again, these mechanisms could work in concert with the physical forces and the associated increase in contact zones. However, we did not observe clear events of lysis in our time-lapse experiments, which make us believe that this factor either had a relatively minor contribution in our experimental system or it remained imperceptible at the densely populated species contact zone. Finally, volatile compounds are also known to play a role in bacterial interactions (Bos et al., 2013; Hou et al., 2021; Netzker et al., 2020). In our experiments, we observed that *P. aeruginosa* colony morphology changed drastically in monoculture when *K. pneumoniae* was grown on neighbouring (yet physically separated) agarose pads (Figure S3). This serendipitous observation suggests that volatiles could influence interactions between pathogens.

In conclusion, our time-lapse microscopy approach allowed us to assess the ecological interactions between *P. aeruginosa* and six other pathogens. We found that interactions are very specific to the species involved and cover a broad spectrum from mutualism to antagonism, suggesting that pathogen interactions are more diverse than previously thought. Important to note is that ecological relationships between two species might change depending on environmental (biotic and abiotic) conditions. Nonetheless, it was surprising to see that *P. aeruginosa* – typically considered a highly competitive species – experienced negative fitness consequences in four out of six cases in our setup. Physical forces due to rapid colony expansion, possibly together with chemical and contact-dependent mechanisms, emerged as important determinants of competitive superiority on agarose pads. A next steps would involve the establishment of a solid framework that allows to predict how different ecological interactions between pathogens affect virulence in polymicrobial infections.

## Methods

### Bacterial strains and growth conditions

We used a fluorescently tagged variant of *Pseudomonas aeruginosa* PAO1 (ATCC 15692) to distinguish it from the other pathogen species referred to as cohabitants. *P. aeruginosa* constitutively expresses the GFP protein which was chromosomally integrated at the neutral attTn7 site in the PAO1 wildtype background (*attTn7::ptac::gfp*). The six cohabitants were all untagged strains and comprised *Acinetobacter baumannii* (DSM30007), *Burkholderia cenocepacia* H111, *Escherichia coli* MG1655, *Enterococcus faecium* (DSM20477), *Klebsiella pneumoniae* (DSM30104), and *Staphylococcus aureus* USA300-FPR3757. For all the experiments, we grew the bacteria in tryptic soy broth (TSB, 21 g/L – Sigma-Aldrich, Buchs SG, Switzerland). The TSB added corresponds to 70% of the dose recommended by the provider. We diluted the medium (henceforth called TSB 70%) to prevent rapid overgrow of colonies on the agarose pads.

### Preparation of agarose pads

To prepare the agarose pads, we followed a previously described method (Weigert and Kümmerli, 2017) with some modifications (regarding the percentage of agarose and overnight cold storage). Briefly, we put two gene frames (1.5 x 1.6 cm – ThermoFisher Scientific) on a standard microscopy slide (76 mm x 26 mm, ThermoScientific) washed with ethanol 70% and dried under a laminar flow. We heated 20 ml of TSB 70% with 1.3% of standard agarose (type LE, BioConcept, Allschwill) in a microwave and pipetted an excess of medium (around 500 µl) into each gene frame chamber. We covered the chambers with another washed microscopy slide and let the medium solidify for 1h at room temperature before storing them overnight at 4°C.

### Sample preparation for pairwise-interaction assays

All pathogen species were inoculated from glycerol stocks and grown at 37°C and 170 rpm (Infors HT multitron standard) with aeration in 1.5 ml of TSB 70% in 24-well plates (Falcon) sealed with parafilm (Parafilm M, Bemis) to avoid evaporation. After overnight incubation, all the strains were re-inoculated in liquid medium to subsequently be collected at the exponential phase. To do so, we adjusted the optical densities (measured at 600 nm, OD_600_) of the overnight cultures to the following values: *P. aeruginosa* (OD_600_ = 0.5), *A. baumannii* (OD_600_ = 3.5), *B. cenocepacia* (OD_600_ = 3.5), *E. coli* (OD_600_ = 0.025), *E. faecium* (OD_600_ = 0.03), *K. pneumoniae* (OD_600_ = 0.003) and *S. aureus* (OD_600_ = 0.1) using a spectrophotometer (U-5100 Hitachi). The variation in OD_600_ adjustment was necessary to account for growth rate difference between species. We re-inoculated 15µl of these cultures into 1.5 ml of fresh TSB 70% in 24-well plates sealed with parafilm and grew them for 6h at 37°C and 170 rpm, to ensure that all species reached their exponential phase (Figure S2). After reaching the exponential phase, we adjusted the OD_600_ of cultures such that a 1:1 volumetric mix result into equal cell ratios. The respective OD_600_ values were determined in a pre-experiment and are as follows: *P. aeruginosa* (OD_600_ = 0.1), *A. baumannii* (OD_600_ = 0.15), *B. cenocepacia* (OD_600_ = 0.2), *E. coli* (OD_600_ = 0.1), *E. faecium* (OD_600_ = 0.15), *K. pneumoniae* (OD_600_ = 0.2) and *S. aureus* (OD_600_ = 0.2). Following adjustments, we diluted all cultures 10-fold in fresh TSB 70% and mixed *P. aeruginosa* cultures with the cultures of all the cohabitants individually in a 1:1 volumetric ratio. These co-cultures together with the monocultures were then used to inoculate the agarose pads as described below.

The microscopy slides holding the agarose pads were removed from the fridge and incubated at room temperature for at least 1h. Under (flame) sterile conditions, we removed the top microscopy slide used as a coverslip by carefully sliding it sideways. We then divided the agarose pads within each gene frame into four smaller pads using a sterile scalpel and introduced channels around all pads to secure oxygen supply during the experiment. We pipetted 1.5 µl of the monocultures onto two different pads and the co-cultures onto the remaining two pads. The same procedure was repeated for the second gene frame on the same microscopy slide with cultures from independent overnight cultures. After evaporation of the droplet (around 2min), we sealed the gene frames with a glass coverslip and covered it with microscopy oil (type F immersion liquid, Leica microsystems). For each pair of species (*P. aeruginosa* versus the six other pathogens), we conducted two independent experiments on two different days.

### Time-lapse fluorescence microscopy experiments

Imaging started right after slide preparation was completed. All microscopy experiments were carried out at the Center for Microscopy and Image Analysis of the University of Zurich (ZMB) with a widefield Olympus ScanR HCS system and the Olympus cellSens software. This microscope was equipped with a motorized Z-drive, a Lumencor SpectraX light engine LED illumination system, a Hamamatsu ORCA-FLASH 4.0 V2 camera system (16-bit grayscale images with a resolution of 2048 x 2048), a CellVivo incubation system and chamber to control the environmental conditions, a FITC SEM fluorescence filter to measure GFP signals (excitation = BP 470 ± 24 nm, emission = BP 515 ± 30 nm, dichroic = 485).

On every agarose pad, we identified three different positions with low numbers of individual cells and with cells of both species being present (for co-species pads). We then imaged these positions every 5 min for 10 h with a PLAPON 60x phase oil objective (NA = 1.42, WD = 0.15 mm) and recorded phase contrast (exposure time 50 ms) and FITC SEM (exposure time 200 ms) images. In one out of 12 experiments, we had to reduce the imaging interval to 10 min due to technical reasons, which had however no impact on bacterial growth and interaction patterns.

### Image processing and analysis

Image processing was conducted with FIJI (Schindelin et al., 2012) using a multi-step process. First, we corrected for drift between images of consecutive time points by aligning frames using an existing open-source script (https://github.com/fiji/Correct_3D_Drift). Second, we segmented individual colonies based on the phase contrast images. The image background was subtracted using the rolling ball algorithm in FIJI (radius = 40 pixels, ca. 4 µm) to correct for uneven illumination and increase the contrast between background and colonies. To smooth over gaps within a colony, we applied a Gaussian filter (sigma = 10 pixels, ca. 1 µm). We used the default automatic threshold function to create a segmentation mask and the resulting regions of interest (ROI). Third, we linked ROIs (tracking the same colony) over time, based on the physical overlap of ROIs between consecutive time points. With this approach, we could exclude colonies that grew from outside into the field of view and were thus not present from the start (no ROI overlap with colonies from previous time points). Fourth, we manually corrected our segmentations. Manual corrections involved separating merging colonies either using the fluorescence signal for co-cultures or by eye for colonies of the same species. We also corrected colonies with inaccurate segmentations. We excluded colonies (i) from the same species that merged early during the time-lapse imaging, (ii) that partially grew out of the field of view, and (iii) with uncorrectable segmentation errors (i.e., when colonies fused and could no longer be distinguished).

Following image processing, we used FIJI to automatically measure colony features. Specifically, we quantified the area of colonies (number of pixels converted to µm^2^) over time as a proxy for fitness. We fitted Gompertz models to the colony growth curves to calculate the maximum growth rate and the area under the growth curves (Figure S1). We further determined the centre of mass of each colony at each time point (defined by x-y coordinates). We used the centre of masses to estimate growth directionality (Dg). Dg was calculated using the formula Dg = De/Da where De (Euclidean distance) represents the distance between the centre of mass of a colony in the first and the last time frame and Da (accumulated distance) represents the sum of all the distances between the center of mass of a colony across successive time frames (Limoli et al., 2019; Niggli et al., 2021). The closer Dg is to 1, the more directional is the growth. We calculated Dg for hourly intervals during the 10h experiments. In addition, we calculated the roundness of colonies at each time point using the standard formula on FIJI: roundness = 4 x area / (π x major_axis^2^). The closer the roundness is to 1, the more circular is the colony. Finally, we identified colonies forming double-layering and measured the area of the double layers by manual segmentations using FIJI.

### Statistical analysis

All statistical analyses were performed with R Studio (version 4.1.1). We used two-way ANOVAs to test whether pathogen fitness (maximum growth rate or area under the curve) differs between species (factor 1) and culturing type (factor 2, mono-versus co-cultures). We further fitted an interaction term to the models (species*culture-type) and included a third term (experimental date without interaction) to account for variation between days. We log10-transformed fitness values to meet the assumption of normally distributed residuals. We consulted diagnostic Q-Q plots and results from the Shapiro-Wilk normality test to verify that residuals were normally distributed. We used Tukey’s HSD *post-hoc* test to extract fitness differences between mono- and co-cultures for each species, which were then used to define the ecological interaction type (from mutualism to competition).

To test whether growth directionality and roundness of colonies differ between mono- and co-cultures, we conducted Welch’s two-sample t-tests for data obtained within hourly intervals. Model residuals were checked for normality and p-values were corrected for multiple testing using the false discovery rate (FDR) method (Benjamini and Hochberg, 1995).

Our gene frame design with four agarose pads enabled us to compare the performance of the two monocultures and the co-cultures that grew at the same time in the same frame (Figure 1). This design is ideal to exclude variation that is due to random (not controllable) factors. However, we made one curious observation. We observed that the roundness of *P. aeruginosa* colonies changed on monoculture pads when *K. pneumoniae* (and only this species) was grown within the same gene frame (Figure S3 & Table S7). One explanation we can offer is that *K. pneumoniae* releases volatiles, which affect *P. aeruginosa* colony morphology on neighbouring pads. While we did not follow up on this phenomenon, we had to adjust the statistical analysis comparing colony roundness. Specifically, we combined all roundness data from *P. aeruginosa* monocultures from experiments without *K. pneumoniae*, and compared this data to the roundness of *P. aeruginosa* colonies in co-cultures with *K. pneumoniae*.

## Acknowledgements

We acknowledge the Center for Microscopy and Image Analysis of the University of Zurich for technical support and maintenance of resources. Figure 1 was created with BioRender.com. This project was supported by a grant from the Swiss National Science Foundation (no. 310030_212266).

## Competing interests

The authors declare no competing interests.

## Data availability statement

The data that support the findings of this study are available from the corresponding author upon request.

## Author contributions

CL and RK designed the study. CL conducted the experiments. CL and TW analysed the data. CL and RK interpreted the data. CL and RK wrote the paper with inputs from TW.

## Notes

### Competing Interest Statement

The authors have declared no competing interest.

