## Supplementary material for "Interactions between *Pseudomonas aeruginosa* and six opportunistic pathogens cover a broad spectrum from mutualism to antagonism": Laffont_SupportingInformation.pdf

Corresponding authors:

Supporting Information contains:

- 3 supplementary figures: Figure S1-S3 (in this file)
- 6 supplementary movies: Movie S1-S6 (as individual movies)
- 7 supplementary tables: Table S1-S7 (in a separate excel file)

**Supplementary figures:**

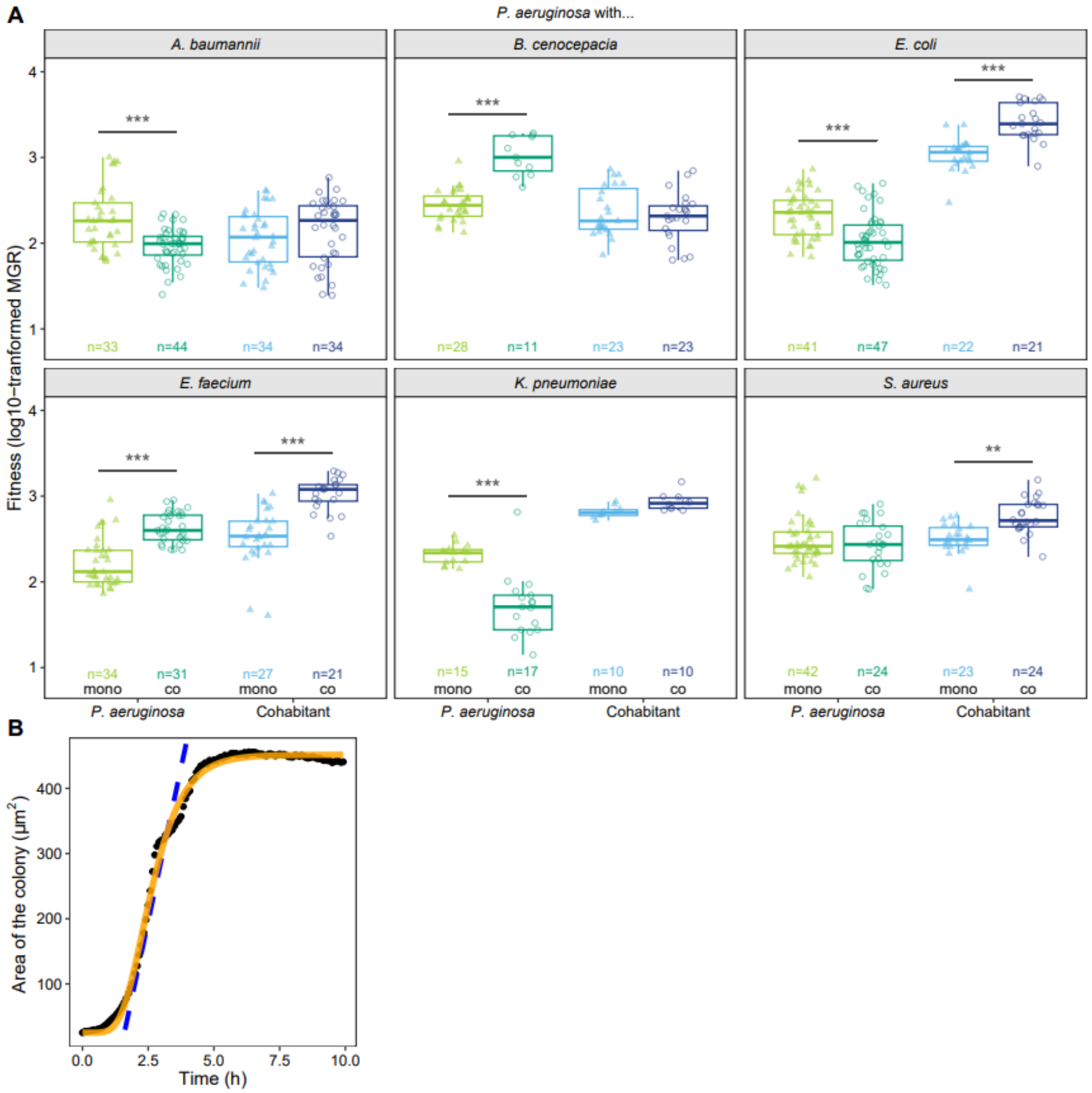


**Figure S1: The maximum growth rate is affected when *P. aeruginosa* is co-cultured with other pathogens. (A)** The boxplots depict the log10-transformed fitness (MGR: maximum growth rate) of *P. aeruginosa* (green) and its cohabitants (blue) in monocultures (light triangles) and co-cultures (dark circles). n-values indicate the total number of colonies tracked for each species combination. Two-way ANOVAs were used in combination with TukeyHSD *post-hoc* tests to examine fitness differences between mono- and co-cultures for *P. aeruginosa* and its cohabitants. Asterisks show the level of significance: * $p_{adj} < 0.05$, ** $p_{adj} < 0.01$, *** $p_{adj} < 0.001$. (B) Gompertz models were used to quantify the maximum growth rate from the growth curves. The black points depict the real data, while the orange curve represents the model fit. The blue dashed line shows the maximum slope of the curve (based on the fitted model) used to calculate the maximum growth rate.

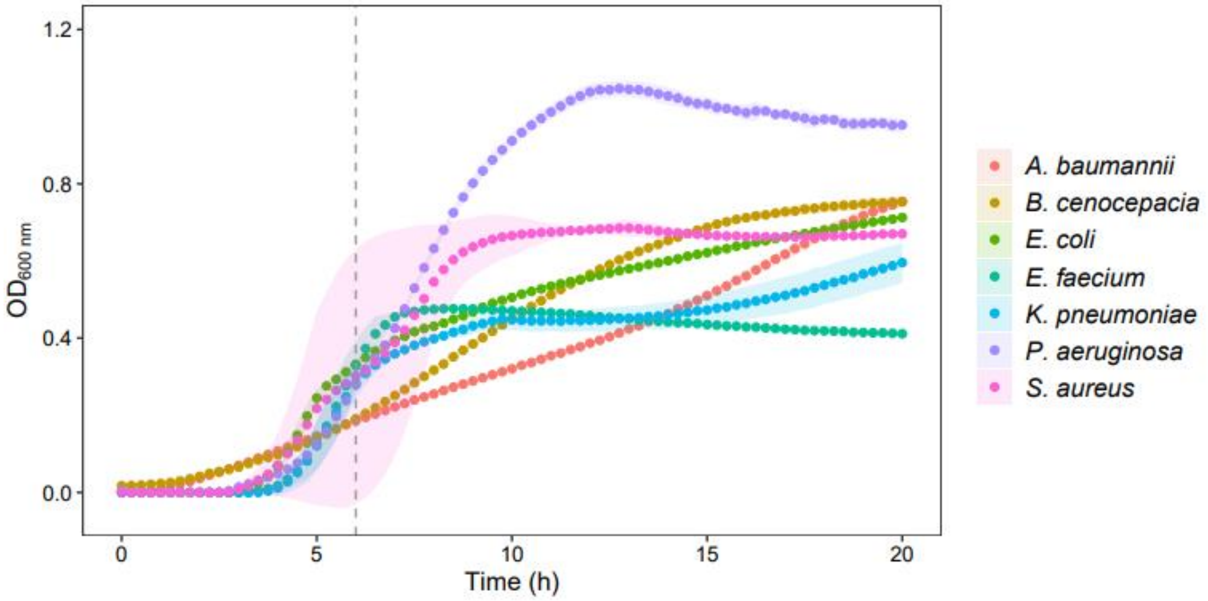


**Figure S2: Cultures of all species were collected at the exponential growth phase to prepare samples for microscopy experiments.** Following growth overnight in TSB, all pathogen species were diluted and re-grown for 6h in fresh medium in order to reach their exponential growth phase. The data represent the means (points) and the 95% confidence intervals (shaded areas) calculated from 3 biological replicates. Growth was tracked in 1.5mL volumes of TSB 70% in 24-well plates, incubated at 37°C in a plate reader (Tecan). Absorbance measurements were taken at 600nm every 15min during 20 hours. Prior to each measurement, cultures were shaken (orbital shaking of 3mm) for 30 seconds.

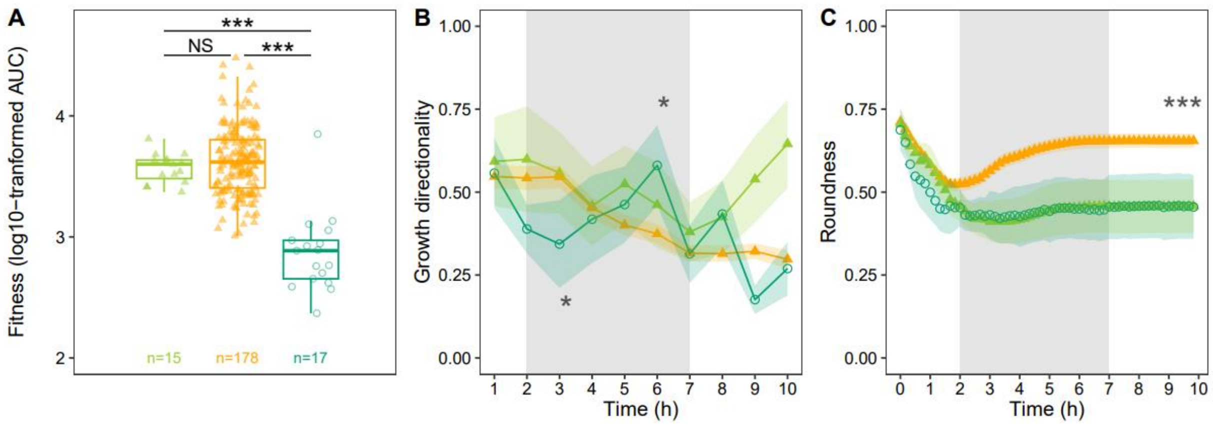


**Figure S3: The roundness of *P. aeruginosa* monocultures colonies is affected by *K. pneumoniae* colonies growing on nearby, yet physically separated, agarose pads.** (A) Fitness, (B) growth directionality and (C) roundness of *P. aeruginosa* colonies grown (i) as monocultures in the absence of *K. pneumoniae* (orange triangles) within the same gene frame, (ii) as monocultures in the presence of *K. pneumoniae* within the same gene frame (light green triangles), and (iii) as co-cultures with *K. pneumoniae* (dark green circles). Data points and shaded areas (B and C) show the means and the 95% confidence intervals across all colonies experiencing the same conditions. The grey shaded areas show the time window (2$^{nd}$ to 7$^{th}$ hour) during which most of the growth occurred. The boxplots (A) depict the log10-transformed values of the AUC (area under the growth curve). Asterisks show the level of significance from the TukeyHSD *post-hoc* tests. NS means non-significant.

The comparisons show that *P. aeruginosa* monoculture fitness is not affected by the absence or presence of *K. pneumoniae* monocultures within the same gene frame (A, light green versus orange triangles). Similarly, there is no difference in the growth directionality during the time of actual growth between the two types of monocultures (B), although there are certain differences (at 3h and 6h) when comparing the two monocultures to the co-cultures. Most importantly, we observed a clear difference between the two *P. aeruginosa* monocultures (i) and (ii) regarding colony roundness (C). Specifically, *P. aeruginosa* colony roundness was high in all experiments without *K. pneumoniae*, whereas *P. aeruginosa* colony roundness dropped in all experiments with *K. pneumoniae*, even when *K. pneumoniae* grew on physically separated agarose pads. This finding suggests that *K. pneumoniae* produces volatiles that impact *P. aeruginosa* colonies on adjacent pads.
